# E3 ligases RNF43 and ZNRF3 display differential specificity for endocytosis of Frizzled family members

**DOI:** 10.1101/2023.11.23.568206

**Authors:** Jeroen M. Bugter, Peter van Kerkhof, Eline Janssen, Thi Tran Ngoc Minh, Daniel Iglesias van Montfort, Cara Jamieson, Ingrid Jordens, Madelon M. Maurice

## Abstract

The transmembrane E3 ligases RNF43 and ZNRF3 perform key tumour suppressor roles by inducing endocytosis of members of the Frizzled (FZD) family, the primary receptors for WNT. Loss-of-function mutations in *RNF43* and *ZNRF3* mediate FZD stabilisation and a WNT-hypersensitive growth state in various cancer types. Strikingly, *RNF43* and *ZNRF3* mutations are differentially distributed across cancer types, raising questions about their functional redundance. Here, we compare the efficacy of RNF43 and ZNRF3 of targeting different FZDs for endocytosis. We find that RNF43 preferentially downregulates FZD1/FZD5/FZD7, while ZNRF3 displays preference towards FZD6/FZD10. We show that the RNF43 transmembrane domain (TMD) is a key molecular determinant for inducing FZD5 endocytosis. Furthermore, a TMD swap between RNF43 and ZNRF3 re-directs their preference for FZD member downregulation. We conclude that RNF43 and ZNRF3 preferentially downregulate specific FZDs by a TMD-dependent mechanism. In accordance, tissue-specific expression patterns of FZD homologues correlate with the incidence of *RNF43* or *ZNRF3* cancer mutations in those tissues. Consequently, our data point to druggable vulnerabilities of specific FZD receptors in *RNF43-* or *ZNRF3*-mutant human cancers.

## Introduction

The closely related transmembrane E3 ligases RNF43 and ZNRF3 control cell surface abundance of Frizzled (FZD) proteins, the primary receptors for WNT, by promoting their ubiquitination and lysosomal degradation (Hao et al., 2012; Koo et al., 2012). Since *RNF43* and *ZNRF3* both represent bona fide WNT target genes, both proteins constitute a negative feedback regulatory role within the WNT cascade (Koo et al., 2012; Takahashi et al., 2014). Within the stem cell niche, this negative feedback role is inactivated by high local concentrations of RSPO proteins, to allow for strong potentiation of WNT signalling required for the maintenance of adult stem cell populations (Hao et al., 2012).

In the mouse small intestine, deletion of either *Rnf43* or *Znrf3* did not induce a discernible phenotype, while the combined deletion of both genes resulted in expansion of the stem cell zone, and eventually adenoma formation (Koo et al., 2012). The essential roles of *Rnf43* and *Znrf3* in adult stem cell homeostasis were shown for multiple tissues, including the liver, adrenal gland and tongue (Basham et al., 2019; Belenguer et al., 2022; Lu et al., 2022). Of note, while both ligases show redundant expression in certain tissues, such as the intestine, in other tissues homeostasis depends on expression of only one of the two ligases. Interestingly, these observations correlate with the differential prevalence of *RNF43* and *ZNRF3* mutations in human cancer subtypes originating from specific tissues (Bugter et al., 2021). The functional relevance of this differential mutational pattern of both E3 ligases remains unclear.

WNT signalling regulation at the cell surface is complex, illustrated by the encoding of 19 WNT genes, 10 FZD genes and various co-receptor genes within the human genome (Niehrs, 2012). Downstream WNT-FZD signalling can be divided in β-catenin-dependent (canonical) and β-catenin-independent (non-canonical) pathways. Certain WNTs or FZD receptors are reported to activate either canonical or non-canonical WNT signalling, while others can activate both pathways depending on the recruitment of co-receptors (Niehrs, 2012). Various studies indicate that RNF43 and ZNRF3 are capable of inhibiting both β-catenin-dependent and -independent WNT pathways (Hao et al., 2012; Moffat et al., 2014; Radaszkiewicz et al., 2021; Tsukiyama et al., 2015). The mechanism by which RNF43/ZNRF3 interact with members of the FZD family however has remained a debated issue. While one study proposed that the extracellular protease associated (PA) domain of RNF43 interacts with the cysteine-rich domain (CRD) of FZD (Tsukiyama et al., 2015), others reported that the RNF43 PA domain is dispensable for WNT signalling suppression (Radaszkiewicz et al., 2020). In another model, intracellular bridging of RNF43 and FZD is mediated by the scaffold protein Dishevelled (DVL) (Jiang et al., 2015), but also these findings have remained disputed due to the observation that RNF43 may interact with FZD in a manner independent of the RNF43-DVL interaction (Tsukiyama et al., 2015, 2020). Importantly, both models do not allow for the identification of receptor specificity. Thus, the precise mechanism of RNF43/ZNRF3-mediated FZD recognition and downregulation remains unaddressed.

WNT hypersensitivity has emerged as a major driver of cancer growth, which may be caused either by mutational inactivation of *RNF43/ZNRF3* or overexpression of RSPO-fusion proteins (Koo et al., 2012; Seshagiri et al., 2012). These observations have led to a growing interest in the application of upstream WNT signalling inhibitors as an anti-cancer strategy (Bugter et al., 2021). As a result, several pan-FZD inhibitors and WNT secretion inhibitors are currently evaluated in clinical trials. Although conclusive clinical efficacy data are still lacking, these strategies are accompanied with side effects caused by a lack of selectivity and/or tissue specificity (Jung & Park, 2020; Zhong & Virshup, 2020). For several cancer subtypes, however, cancer growth was shown to rely on one specific FZD homologue, which holds promise for more selective inhibition strategies (Do et al., 2022; Steinhart et al., 2017).

The level of functional redundancy of RNF43- and ZNRF3-mediated FZD targeting has remained unclear. Over recent years, several studies reported unique roles of post-translational regulation of one of the two ligases, or unique roles and activities beyond WNT signalling (Ci et al., 2018; Fang et al., 2022; Kim et al., 2021; Spit et al., 2020). Furthermore, the observation that several cancer types display specific loss of only one of the two ligases suggests that non-redundant functions exist. In this study, we uncover specific roles of RNF43 and ZNRF3 in the downregulation of FZD receptors. In addition, by examining tissue-specific expression patterns we translate FZD-specific roles to the incidence of *RNF43* and *ZNRF3* mutations in cancer.

## Results and Discussion

### RNF43 and ZNRF3 suppress WNT3A- and WNT5A-mediated signalling

To examine the activity of overexpressed RNF43 or ZNRF3 variants and avoid effects of endogenous protein expression, we generated *RNF43/ZNRF3* double knockout HEK293T cells (R/ZdKO). Similar to previous reports (Jiang et al., 2015; Radaszkiewicz et al., 2020; Spit et al., 2020), R/ZdKO cells displayed a strong increase in basal WNT/β-catenin signalling activity in comparison to wild type (WT) HEK293T cells, which was further enhanced by WNT3A stimulation, as shown by a WNT luciferase reporter assay (Fig. 1A). As expected, R/ZdKO cells were unresponsive to RSPO1 treatment (Fig. 1A) (Jiang et al., 2015; Radaszkiewicz et al., 2020). For evaluation of levels of β-catenin-independent WNT signalling induced by WNT5a (González-Sancho et al., 2004; Yamaguchi et al., 1999), we examined phosphorylation levels of DVL2, an immediate downstream effector of FZD-mediated signalling that acts in both β-catenin-dependent and -independent WNT pathways (Bryja et al., 2007; González-Sancho et al., 2004). We blocked interference of endogenous WNT ligands using WNT-secretion inhibitor C59. R/ZdKO cells displayed increased levels of DVL2 phosphorylation upon WNT3A or WNT5A stimulation in comparison to WT cells (Fig. 1B). Co-treatment with RSPO1 further increased the levels of DVL2 phosphorylation of WT cells, while R/ZdKO cells were insensitive to RSPO1 treatment (Fig. 1B). Overexpression of either RNF43 or ZNRF3 in R/ZdKO cells reduced levels of DVL2 phosphorylation upon WNT3A or WNT5A treatment (Fig. 1C). In line with previous findings, these results suggest that both RNF43 and ZNRF3 possess the capacity to inhibit both β-catenin-dependent and -independent WNT pathways (Hao et al., 2012; Moffat et al., 2014; Radaszkiewicz et al., 2021; Tsukiyama et al., 2015).

**Figure 1.**
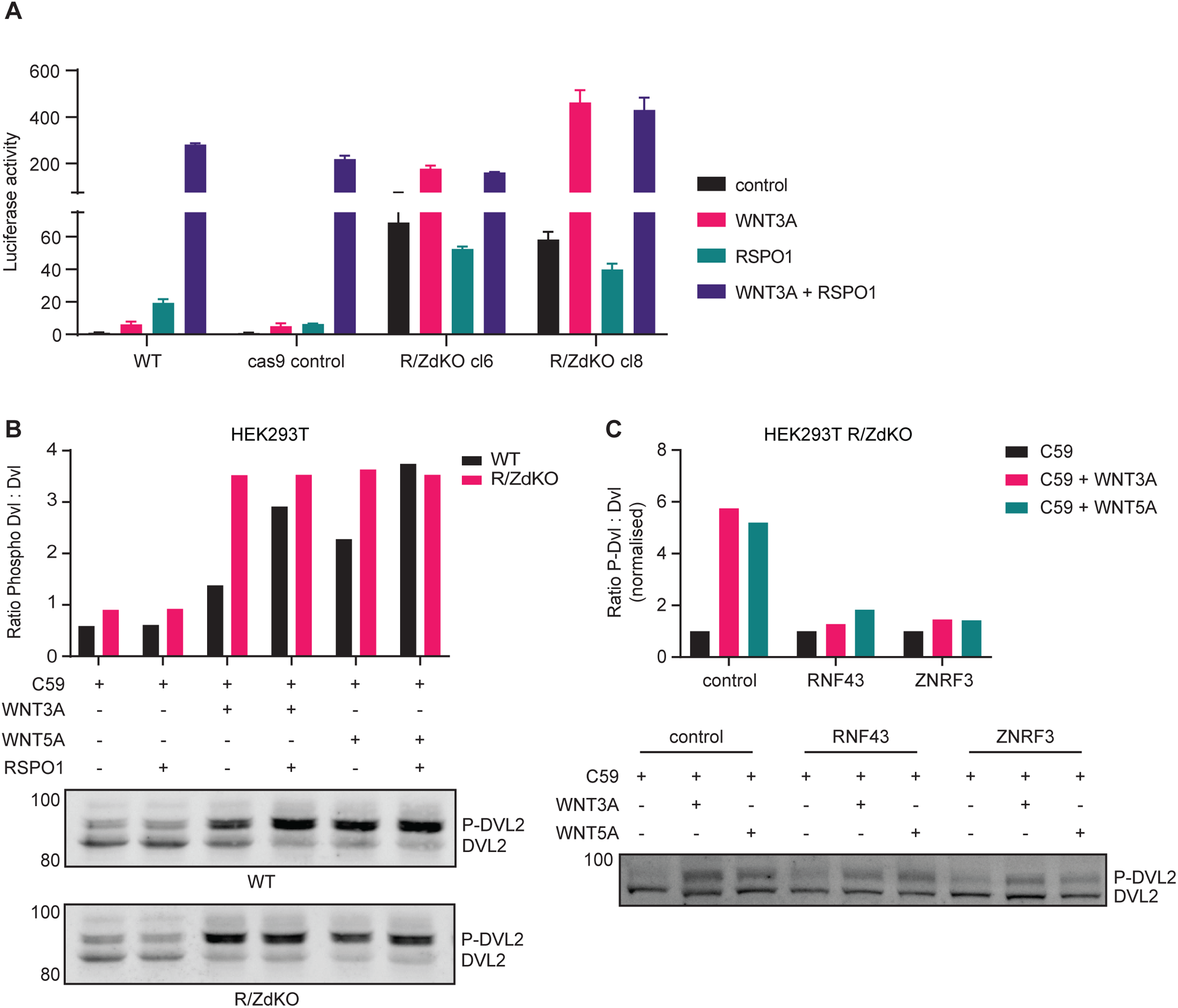
RNF43 and ZNRF3-mediated suppression of WNT3A- and WNT5A-induced signalling. **A.** WNT/β-catenin-mediated luciferase reporter assay performed with wildtype (WT) HEK293T cells, HEK293T cas9-expressing control cells or two clonal HEK293T RNF43/ZNRF3 double knockout (R/ZdKO) cell lines that were stimulated with WNT3A and/or RSPO1-conditioned medium (CM) compared to control medium. **B.** Western blot analysis of DVL2 phosphorylation in WT HEK293T and R/ZdKO cells after stimulation with recombinant WNT3A or WNT5A and/or RSPO1-CM. All samples were treated with Porcupine inhibitor (C59) to exclude effects of endogenous WNTs. Ratio between phosphorylated DVL2 (P-DVL2) and unphosphorylated DVL2 protein band density was calculated and plotted. A representative experiment (N=4) is shown. **C.** Western blot analysis of DVL2 phosphorylation in HEK293T R/ZdKO cells overexpressing RNF43, ZNRF3 or control vector after stimulation with recombinant WNT3A or WNT5A.

### RNF43 and ZNRF3 preferentially target specific FZD receptors for endocytosis

Based on their differential tissue expression and mutational patterns (Bugter et al., 2021), we wondered if RNF43 and ZNRF3 may display a preference for targeting specific FZDs for endocytosis (Hao et al., 2012; Koo et al., 2012). To investigate this issue, we employed a set of expression constructs for nine human FZDs (Do et al., 2022)and examined their sensitivity for ligase-induced downregulation from the plasma membrane.

For most FZDs, except for FZD4, we observed a clear distinction between higher and lower molecular weight (MW) bands on Western blot (Fig. 2A). Previously, for FZD5 we determined that the higher MW band represents the mature, post-Golgi glycosylated form, while the lower band represents an immature high mannose-modified ER-localised form (Koo et al., 2012). Using cell surface biotinylation and streptavidin pull-downs, we confirmed that for all FZD members displaying two MW forms, the higher molecular weight form represents the mature, plasma membrane-localised fraction of the protein (Fig. S1). FZD6 and FZD8 showed relatively low expression levels and for those homologues the mature protein forms were detectable only after immunoprecipitation (Fig. 2A, S1A). Of note, FZD3 did not express at all and was therefore excluded from our analysis.

**Figure 2.**
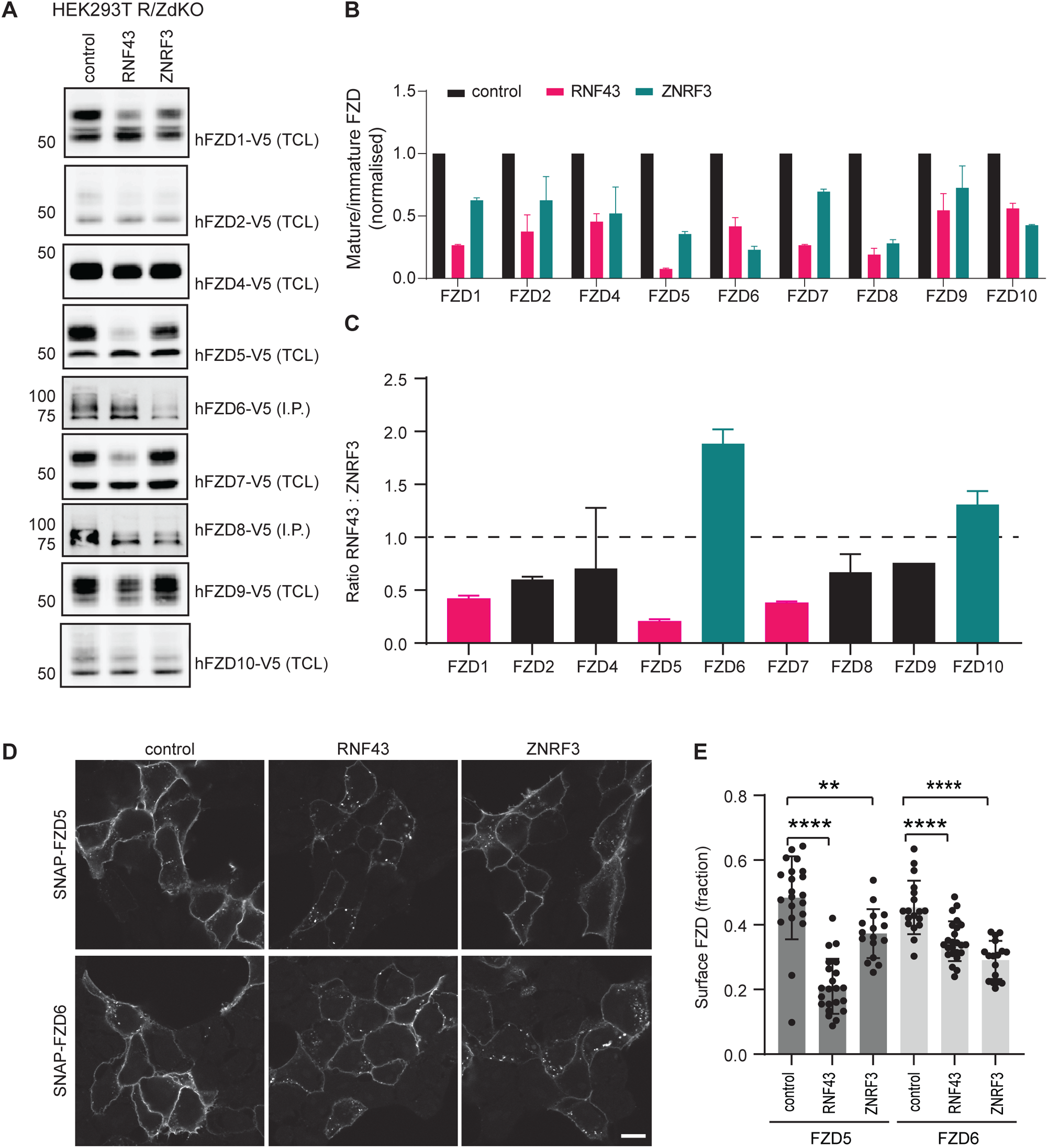
RNF43 and ZNRF3 display Frizzled substrate specificity. **A.** Western blot analysis of V5-tagged human FZD constructs upon co-expression with RNF43, ZNRF3 or control vector. V5-immunoprecipitated (IP) samples are blotted in case of FZD6 and FZD8, total cell lysate (TCL) is blotted in case of other FZDs. **B.** Relative levels of mature FZD are shown as the ratio between mature and immature FZD forms, normalised to pcDNA3-transfected control. N=3. **C.** Ratio between RNF43 and ZNRF3 downregulation efficacy for each FZD. Green = ZNRF3-preferred substrate, Pink = RNF43-preferred substrate. N=3. **D.** Immunofluorescence showing the subcellular localisation of SNAP–FZD5 or SNAP-FZD6 co-transfected with RNF43, ZNRF3 or control vector. Surface SNAP–FZD was labelled for 15 min and chased for either 30 (FZD5) or 60 min (FZD6). Scale bar, 10 µm. **E.** Quantification of surface FZD fractions as shown in (D). Graph shows a representative experiment with the average surface fractions for the indicated conditions (N=16-24 cells). The statistical significance was determined by a one-way Anova and is indicated as **** (p<0.0001) or ** (p<0.01).

We co-expressed each FZD with RNF43 or ZNRF3 in R/ZdKO cells and examined the capacity of the two ligases to degrade individual FZD members, by calculating the mature-to-immature ratio of FZD by Western blotting (Fig. 2A, B). Strikingly, both ligases displayed a clear preference for downregulating a specific subset of FZDs, in a partially overlapping pattern. Notably, RNF43 most potently downregulated FZD1, FZD5 and FZD7, while ZNRF3 displayed a preference for FZD6 and FZD10 (Fig. 2C). To further examine ligase specificity for these FZD subgroups, we co-expressed RNF43 or ZNRF3 with FZD5 or FZD6 containing an extracellular SNAP-tag and performed a pulse chase experiment to determine the level of ligase-mediated FZD internalisation. Similar to our previous findings (Koo et al., 2012), overexpression of RNF43 induced rapid reduction of the FZD5 pool that resides at the cell surface (Fig. 2D,E, Fig. S2). ZNRF3, on the other hand, only caused a moderate reduction of the surface fraction of FZD5. Opposite effects were observed for FZD6, with ZNRF3 inducing a more efficient internalisation as compared to RNF43 (Fig. 2D,E). Thus, although RNF43 and ZNRF3 both suppress WNT signalling via FZD downregulation, each homologue possesses a ligase-specific substrate preference.

### Molecular requirements for RNF43-mediated FZD downregulation

To determine the protein regions that confer substrate specificity of RNF43 and ZNRF3, we first replaced their endogenous signal peptides by the highly efficient H2-Kb signal peptide commonly used for optimized expression of type I membrane proteins (Bénaroch et al., 1995; Fiebiger et al., 2005). Replacement of the signal peptide did not affect the capacity of RNF43 and ZNRF3 to differentially downregulate FZD5 (Fig. 3A), indicating that substrate specificity must be dictated by different regions within these ligases.

**Figure 3.**
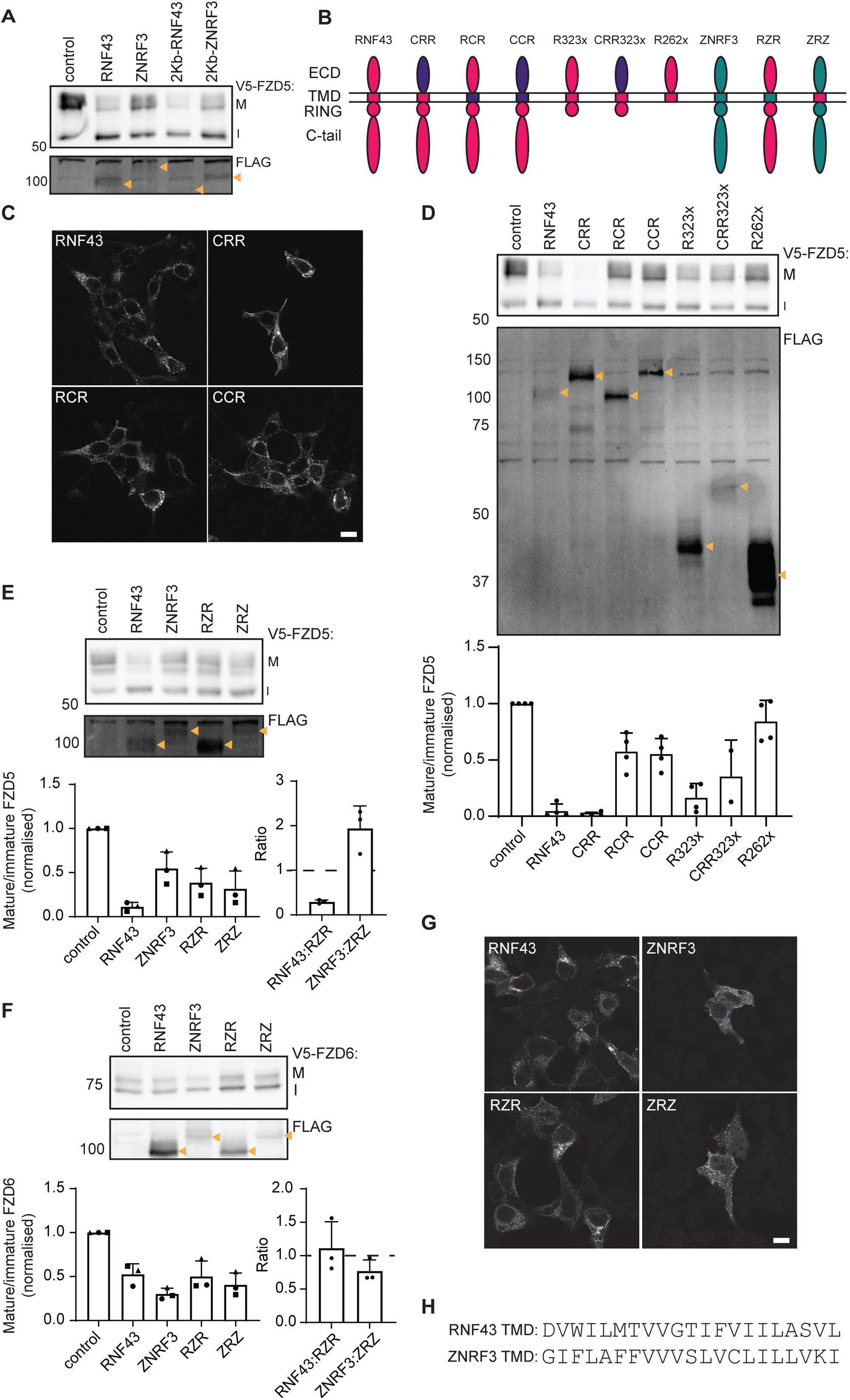
The RNF43 transmembrane domain is essential for FZD downregulation and determines FZD specificity. **A.** FZD5 downregulation capacity of RNF43 and ZNRF3 with their endogenous signal peptides or a generic H-2Kb signal peptide. Mature (M) and immature (I) FZD5 are indicated. Expression of FLAG-tagged ligases is indicated with yellow arrows. **B.** Schematic representation of the RNF43 (pink), ZNRF3 (green) and CD16/7 (purple) chimeric and truncated constructs used in this study. Extracellular domain (ECD), transmembrane domain (TMD), E3 ligase RING domain (RING), C-terminal tail (C-tail). **C.** Immunofluorescence showing the subcellular localisation of RNF43, CRR, RCR and CCR in HEK293T cells. Scale bar, 10 µm. **D.** FZD5 downregulation capacity of RNF43 constructs with various protein domains replaced or truncated. Blots are labelled as (A). Relative levels of mature FZD5 are shown as the ratio between mature (M) and immature (I) forms, normalised to pcDNA3-transfected control. N=4. **E,F.** FZD5 **(E)** and FZD6 **(F)** downregulation capacity of RNF43, ZNRF3 and transmembrane domain (TMD)-swapped RZR and ZRZ constructs. Blots are labelled as in (A). Relative levels of mature FZD are shown as in (D) for indicated constructs (with or without swapped TMD). Ratios between FZD levels upon RNF43 and RZR or ZNRF3 and ZRZ co-expression are plotted as well. N=3. **G.** Immunofluorescence showing the subcellular localisation of RNF43, ZNRF3, RZR and ZRZ in HEK293T cells. Scale bar, 10 µm. **H.** Comparison of amino acid sequence of the transmembrane domains of RNF43 and ZNRF3 as described on Uniprot.

We next investigated which regions of the RNF43 protein are required for FZD5 downregulation, by generating various RNF43 deletion and chimeric variants. We replaced the extracellular domain (ECD) of RNF43 by the extracellular domain of the unrelated protein CD16, and we swapped its transmembrane domain (TMD) for the similarly sized TMD of CD7 (Fig. 3B). We confirmed that the subcellular localisation of each of these variants was comparable with WT RNF43 (Fig. 3C). Functionally, we found that replacement of RNF43-ECD with CD16-ECD (CRR) did not affect the capacity of RNF43 to downregulate FZD5 (Fig. 3D). These findings indicate that RNF43-ECD is dispensable for FZD5 downregulation and suppression of WNT/β-catenin signalling, in agreement with previous findings (Radaszkiewicz et al., 2020), but in disagreement with another report (Tsukiyama et al., 2015). Strikingly, replacement of RNF43-TMD with CD7-TMD significantly diminished the capacity of RNF43 to downregulate FZD5 (Fig. 3D). These findings thus reveal a previously unknown role of RNF43-TMD for efficient FZD5 downregulation.

In an earlier report, we showed that a large fraction of the intracellular C-terminus of RNF43 is dispensable for its role in FZD5 downregulation (Spit et al., 2020). To further map the functionally required region, we truncated the RNF43 C-terminus further, removing all known functional domains downstream of the RING-domain (variant R323x; Fig. 3D). Functional analysis revealed that the whole cytosolic tail downstream of the RING domain is dispensable for RNF43-mediated downregulation of FZD5 (Fig. 3D). Notably, this functionally competent RNF43 truncant also lacks the previously identified DVL Interaction Region (DIR), which was proposed to be essential for the interaction of RNF43 with FZD (Jiang et al., 2015). Even a chimeric RNF43 variant carrying a CD16-derived ECD along with an early truncation downstream of the RING domain (R323X) retained functionality in these assays, albeit with somewhat attenuated activity (Fig. 3D). Thus, our results indicate that the RNF43-TMD and RING domain comprise the minimally required fraction of the protein for FZD5 downregulation.

### The transmembrane domain determines RNF43 and ZNRF3 substrate specificity

We next wondered whether substrate specificity of RNF43 and ZNRF3 is dictated by their TMD. To address this issue, we swapped the transmembrane domains of the two ligases (Fig. 3B) and tested their efficacy in downregulating FZD5 and FZD6. Strikingly, ZNRF3 containing the RNF43 TMD showed an increased FZD5 downregulation efficacy compared to WT ZNRF3, whereas RNF43 containing the ZNRF3 TMD showed a decreased efficacy in downregulating FZD5 compared to WT RNF43 (Fig. 3E). By contrast, ZNRF3 carrying the RNF43-TMD showed a reduced capacity to downregulate FZD6 compared to WT ZRNF3, whereas the capacity of RNF43 to downregulate FZD6 remained unaltered following the TMD swap (Fig. 3F) Importantly, swapping of TMDs did not affect the overall intracellular localisation of both ligase variants (Fig. 3G). These data indicate that the substrate specificity of these closely related E3 ligase proteins is dictated, at least partly, by their transmembrane domains.

### Relation between tissue distribution of RNF43 and ZNRF3 cancer mutations and Frizzled expression

To assess the biological relevance of our findings, we examined the distribution of FZD expression across various human tissue types. Remarkably, FZD expression patterns displayed a substantial level of tissue specificity. While some tissues showed higher expression of RNF43-preferred substrates (FZD1, 5, 7) others mainly expressed ZNRF3-preferred substrates (FZD6, 10) (Fig. 4A, S3). FZD5 is the dominant homologue in pancreas, colon, small intestine and liver. By contrast, FZD6 is the dominant homologue in adrenal gland and skin (Fig. 4A). Notably, cancer types arising from the latter two organs are enriched for mutations in *ZNRF3* in comparison to mutations in *RNF43* (Fig. 4B). In line with these observations, *Znrf3* loss, but not *Rnf43* loss, is sufficient to transform adrenal cortical tissue in mouse models (Fig. 4C) (Basham et al., 2019), although both ligases are expressed in these tissues (Fig. S3). By contrast, *RNF43* mutations, but not *ZNRF3* mutations, are enriched in pancreatic ductal adenocarcinoma (PDAC) (Fig. 4B), and *Rnf43* loss promotes PDAC formation in *Kras*-mutant mouse models (Fig. 4C) (Hosein et al., 2022). Accordingly, *RNF43*-mutant PDAC cell lines and xenografts appear exclusively dependent on signalling via FZD5 (Steinhart et al., 2017). Another example is provided by the critical role of RNF43 but not ZNRF3 in oligodendrocyte maturation upon injury (Niu et al., 2021). This process requires the downregulation of FZD1 that we identified as an RNF43-preferred substrate.

**Figure 4.**
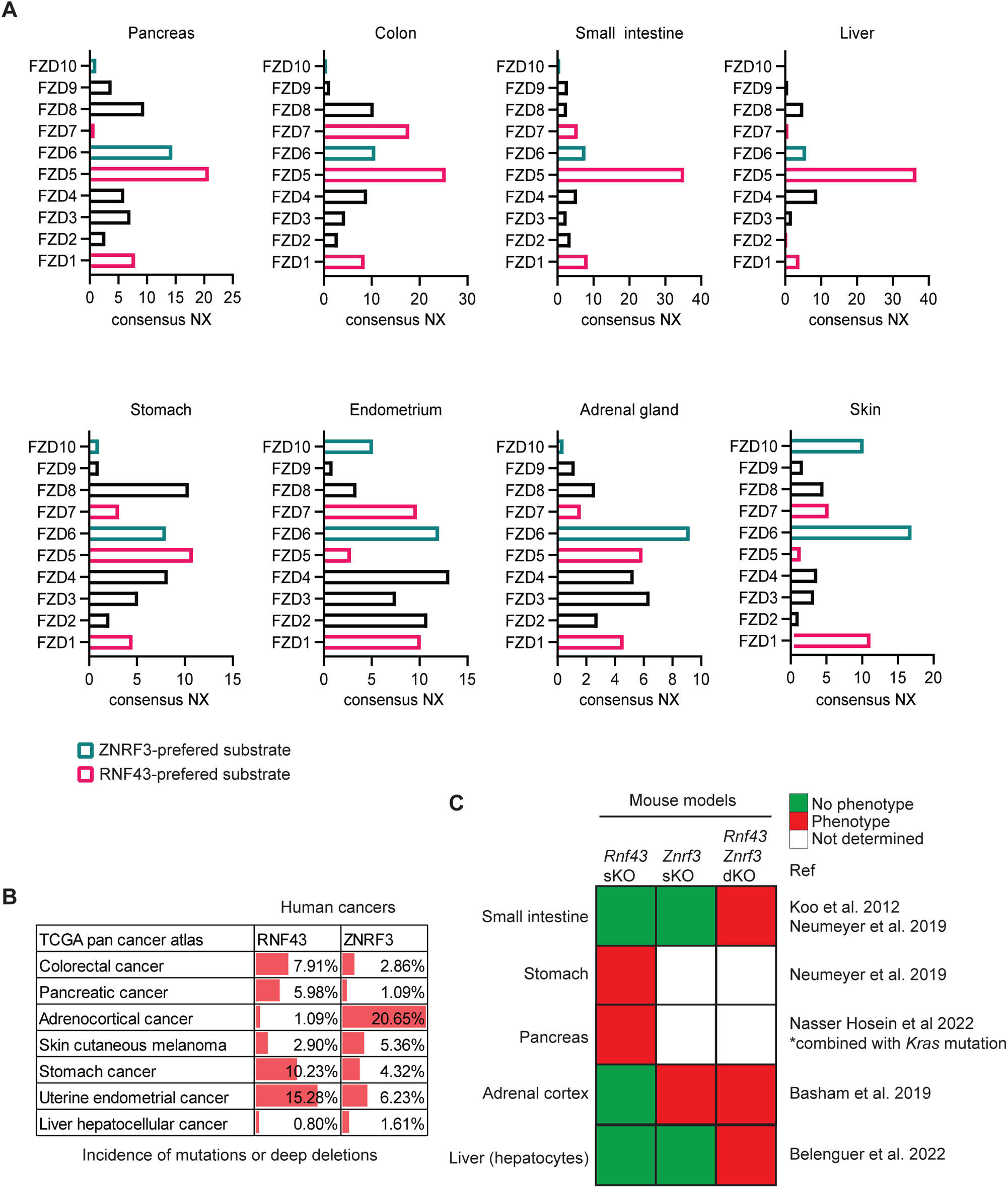
Frizzled homologue expression in various human tissues relates to incidence of *RNF43* and/or *ZNRF3* mutations in cancer or phenotypes in mouse models. **A.** Consensus normalised expression (NX) of all FZD genes among human tissues. RNF43- and ZNRF3-preferred substrates are indicated in pink and green, respectively **B.** Incidence of RNF43 and ZNRF3 mutations and deep deletions among TCGA pan cancer atlas datasets. **C.** Results of published mouse studies indicated reported phenotype of *Rnf43*, *Znrf3* single-knockout (sKO) or double-knockout (dKO).

In conclusion, we here reveal a substantial level of substrate specificity for the closely related E3 ligases RNF43 and ZNRF3 that were previously considered as functionally redundant (Koo et al., 2012). Furthermore, our results uncover a role of the TMD of these ligases in determining substrate preference. In line with these findings, the amino acid sequence of the TMD is poorly conserved between RNF43 and ZNRF3, with only 1/21 (4,7%) overlap in amino acid composition (Fig. 3H). By contrast, the PA domain displays 46/100 (46%) and the RING domain 31/45 (69%) overlap in amino acids for both proteins (Tsukiyama et al., 2021). Regulation of FZD activity was shown previously to depend on TMD-mediated interactions. For instance, the TMD domain of TMEM59 was shown to promote clustering of FZD receptors (Gerlach et al., 2018), and dimerisation of FZD6 is dependent on homotypic TMD-TMD interactions (Petersen et al., 2017). Whether the TMDs of RNF43 and ZNRF3 are involved in direct intramembrane interactions with FZDs or rather affect their localisation to specific membrane domains remain open questions for further investigation. Overall, our findings provide an explanation for the differential incidence of tissue-specific mutation frequency of *RNF43* and *ZNRF3* in cancers. Understanding the preferences of these highly mutated E3 ligases might help in predicting vulnerabilities to inhibition of specific FZD receptors in *RNF43-* or *ZNRF3*-mutant cancers.

## Acknowledgements

We thank Karl Willert (Department of Cellular & Molecular Medicine, University of California San Diego, San Diego, United States) for kindly providing the library of FZD(1-10)-V5-IRES-mKate constructs. We thank members of the laboratory of M.M.M. for discussions and suggestions. This work is part of the Oncode Institute, which is partly financed by the Dutch Cancer Society. This work was supported by the ZonMW TOP Grant 91218050 (to M.M.M), Dutch Cancer Society grant 13112 (to M.M.M.) and NWO Gravitation project IMAGINE! (to M.M.M.).

Current addresses: J.M.B.: Institute of Molecular Oncology and Functional Genomics, School of Medicine, Technische Universität München, Munich, Germany; Center for Translational Cancer Research (TranslaTUM), School of Medicine, Technische Universität München, Munich, Germany. T.T.N.M.: Division Cell Biology, Metabolism & Cancer, Department Biomolecular Health Sciences, Faculty of Veterinary Medicine and Biomolecular Mass Spectrometry and Proteomics, Bijvoet Centre for Biomolecular Research, Utrecht University, Utrecht, The Netherlands.

E.J.: Department of Medical BioSciences, Radboud university medical center, Geert Grooteplein Zuid 26-28, 6525 GA Nijmegen, The Netherlands

## Author contributions

JMB, IJ, and MMM conceived and designed the experiments. JMB, PvK, EJ, TTNM, DIvM, CJ, and IJ performed the experiments. JMB, PvK, EJ, TTNM, DIvM, CJ, IJ, and MMM analyzed the data. JMB and MMM wrote the manuscript, which was reviewed by all authors.

## Materials and methods

### Cell culture and generation of knockout lines

Human Embryonic Kidney (HEK) 293T cells were cultured in DMEM high glucose (Invitrogen), supplemented with 10% fetal bovine serum (GE Healthcare), 2 mM UltraGlutamine (Lonza), 100 units/mL penicillin, and 100 μg/mL streptomycin (Invitrogen). Cells were cultured at 37°C in 5% CO2. WNT3a-conditioned medium (CM) and RSPO1-CM were produced as described (Fenderico et al., 2019; Tauriello et al., 2010) Transfections were performed using either FuGENE 6 (Promega) according to manufacturer’s protocol for β-catenin-mediated reporter assays and microscopy or polyethylenimine (PEI) for Western blot analysis.

RNF43 and ZNRF3 double knockout cells (R/ZdKO) were generated by simultaneously targeting *RNF43* (gRNA: ATTGCACAGGTACAGCGGGT) and *ZNRF3* (gRNA: GCCAAGCGAGCAGTACAGCG) with gRNAs cloned in a pSpCas9(BB)-2A-Puro vector (Addgene # 48139). Monoclonal cell lines that were homozygous knockout for both genes were confirmed by genotyping and functional analysis in a β-catenin-mediated reporter assay (Fig. 1a).

### Plasmids and transfection

RNF43–2xFlag–HA (FFHH) and ZNRF3-FFHH constructs and RING mutants were described previously (Koo et al., 2012). V5-mouse FZD5 was described earlier (Tauriello et al., 2012) and V5-human FZD6 was generated similarly. RNF43 mutants were generated by PCR-subcloning using Q5 High-Fidelity 2× Master Mix (NEB). Domain swapped constructs were generated by in-fusion cloning (Takara Bio). FZD(1-10)-V5-IRES-mKate constructs were a kind gift of Karl Willert (Department of Cellular & Molecular Medicine, University of California San Diego, San Diego, United States). All constructs were sequence verified.

Transfections were performed using either FuGENE6 (Promega) according to manufacturer’s protocol for β-catenin-mediated reporter assays and microscopy or polyethylenimine (PEI) for Western blot analysis.

#### β-catenin-mediated reporter assays

HEK293T WT or R/ZdKO cells were seeded in 24-well plates and transfected the next day with 30 ng of reporter construct TOPFlash or FOPFlash, 5 ng of thymidine kinase (TK)-Renilla and the indicated constructs. Cells were stimulated 6 h post-transfection with WNT3a-CM and/or RSPO1-CM overnight, then cells were lysed in Passive lysis buffer (Promega) for 20 min at room temperature. Levels of Firefly and Renilla luciferase were measured using the dual-luciferase kit (Promega) accordingly to the manufacturer’s instructions on a Berthold luminometer Centro LB960.

### WNT3A and WNT5A response assays

Response to canonical and non-canonical WNT ligands, respectively WNT3A and WNT5A, was measured as previously reported (Radaszkiewicz et al., 2021). HEK293T R/ZdKO cells were treated with 10 μM PORCN inhibitor C59 (Tocris) to block secretion of endogenous WNT ligands. Cells were transfected with RNF43, ZNRF3 or control vector and incubated with C59 overnight. The next day, cells were stimulated with 100 ng/mL recombinant WNT3A (R&D Systems, 5036-WN) or WNT5A (R&D Systems, 645-WN). Cells were lysed in RIPA buffer (1% IGEPAL CA-630 + 1% Sodium Deoxycholate + 0,1% SDS + 50 mM NaF + 150 mM NaCl + 25 mM Tris pH 7,5 + 1 mM EDTA + 1 mM PMSF + 10 µg/mL Aprotinin + 10 µg/mL Leupeptin + 1 mM Na_3_Vo_4_) and boiled in 1x SDS sample buffer for 5 min.

### FZD downregulation assays

HEK293T R/ZdKO cells were co-transfected with V5-tagged FZD and RNF43-FFHH, ZNRF3-FFHH or empty vector in 1:1 ratio. The next day, cells were subjected to cell surface biotinylation (see below) or directly lysed in lysis buffer (1% Triton-X100 + 150 mM NaCl + 1 mM EDTA in 20 mM Tris pH 7,5 + 50 mM NaF + 1 mM PMSF + 10 µg/mL Aprotinin + 10 µg/mL Leupeptin + 1 mM Na_3_VO_4_). In case of FZD6 and FZD8, due to low expression, immunoprecipitation was used to concentrate protein levels (see below). In the other cases total protein lysates were eluted in SDS sample buffer and incubated for 30 min at 37°C.

### FZD immunoprecipitation

Protein lysates were incubated with 1 µg mouse anti-V5 antibody (Genscript A01724) for 2 h at 4°C, followed by 1 h incubation with 25 µL Protein A-agarose beads (RepliGen). Beads were washed five times with 0,1 x PBS after which beads were eluted in SDS sample buffer and incubated for 30 min at 37°C.

### Cell surface biotinylation

Live cells were washed three times with ice-cold PBS complete (including Ca^++^ and Mg^++^) and incubated for 30 minutes on ice in PBS complete with 0,8 mM Sulfo-NHS-SS-Biotin (Thermo Fisher Scientific). Cells were washed once with PBS complete, once with DMEM medium + 50 mM Glycine to quench unreacted biotin for 5 min on ice and once with PBS complete. Cells were collected and lysed in lysis buffer. Protein lysates were incubated with 25 µL Streptavidin-agarose beads (Pierce, Thermo Fisher Scientific) for 1 h at 4°C. Beads were washed five times with 0,1 x PBS after which beads were eluted in SDS sample buffer and incubated for 30 min at 37°C.

### Western blotting and analysis

Western blotting was performed using standard procedures. Samples were resolved by SDS-PAGE, transferred to Immobilon-FL PVDF membranes (Milipore, Sigma), blocked with Odyssey blocking buffer (Li-Cor), incubated with the indicated primary antibodies overnight at 4°C, with secondary antibodies for 1 h at room temperature, and imaged using the Amersham Typhoon NIR laser scanner (GE Healthcare). Quantifications were performed using ImageQuant TL 8.2 (GE healthcare). Activation of downstream signalling in the WNT3A and WNT5A response assays was determined by calculating the ratio of phosphorylated over non-phosphorylated DVL2 bands on Western blot (Radaszkiewicz et al., 2021). FZD downregulation was determined by calculating the ratio of the mature over the immature band, except for FZD4 for which the two isoforms were indistinguishable and the total levels were calculated instead.

### Immunofluorescence and SNAP surface labelling

HEK293T cells were seeded on laminin-coated glass coverslips. The next day cells were transfected with indicated constructs. 20 hours post-transfection, cells were fixed in 4% paraformaldehyde for 30 minutes and subsequently the reaction was quenched by incubation with 0,05M NH_4_Cl for 30 minutes at room temperature. Cells were blocked in blocking buffer (PBS with 2% bovine serum albumin (Sigma) and 0,1% saponin). Next, cells were incubated with primary and secondary antibodies diluted in blocking buffer for 1 h at room temperature. Cells were mounted in ProLong Gold (Life Technologies) and analyzed using a Zeiss LSM700 confocal microscope.

For SNAP surface labelling, cells were incubated with 1 μM SNAP–Alexa^488^ (Bioke) for 15 min at room temperature. Subsequently, cells were chased for 30 or 60 min at 37° C and washed once with warm medium before fixation and staining (see above).

### Quantification surface fraction SNAP-FZD

Confocal images were processed in ImageJ (1.53) to determine the surface FZD fractions. To measure the total intensity of FZD per cell, a region of interest (ROI) was drawn lining the outside of the cell. Subsequently, by drawing a ROI right below the cell surface, the intracellular intensity of FZD signal was measured. Next, the surface fraction was calculated by subtracting the intracellular intensity from the total intensity. Next, the fraction of surface levels over total was calculated. The ROI for total FZD was transferred to the second channel to determine the total level of RNF43 or ZNRF3 by anti-Flag labelling (Fig S2). The *p*-values for the differences in FZD surface fraction between control and either RNF43 or ZNRF3 expressing cells were calculated using a one-way Anova using GraphPad Prism 9.5.1.

### Antibodies

| Name | Manufacturer and catalogue number | Dilution for western blot | Dilution for immunofluorescence |
| --- | --- | --- | --- |
| Rabbit anti DVL2 | Cell Signaling, 3216S | 1:1.000 | X |
| Rabbit anti V5 | Sigma, V8137 | 1:2.000 | X |
| Mouse anti FLAG | Sigma, F3165 | 1:5.000 | 1:1000 |
| Rabbit anti FLAG | Sigma, F7425 | 1:2.500 | 1:500 |
| Mouse anti Actin | MP, 691001 | 1:10.000 | x |
| Goat anti Rabbit IRDye680 | Li-Cor, 926-32221 | 1:8.000 | X |
| Goat anti Mouse IRDye680 | Li-Cor, 925-68070 | 1:8.000 | X |
| Goat anti Rabbit IRDye800 | Li-Cor, 926-32211 | 1:8.000 | X |
| Goat anti Mouse IRDye800 | Li-Cor, 926-32210 | 1:8.000 | X |
| Goat anti Rabbit Alexa488 | Invitrogen, | X | 1:300 |
| Goat anti mouse Alexa569 | Invitrogen, | X | 1:300 |

### Public data analysis

To address FZD expression across human tissues, normalised consensus RNA expression data (NX) were downloaded from the Human Protein Atlas version 22.0.

The frequency of *RNF43* and *ZNRF3* mutations in human cancer subtypes was addressed in the cancer genomics atlas (TCGA) Pan Cancer Atlas data that were accessed via Memorial Sloan Kettering Cancer Center (MSKCC) cBioPortal. Mutations and deep deletions were combined.

## Supplementary figures legends

**Figure S1.**
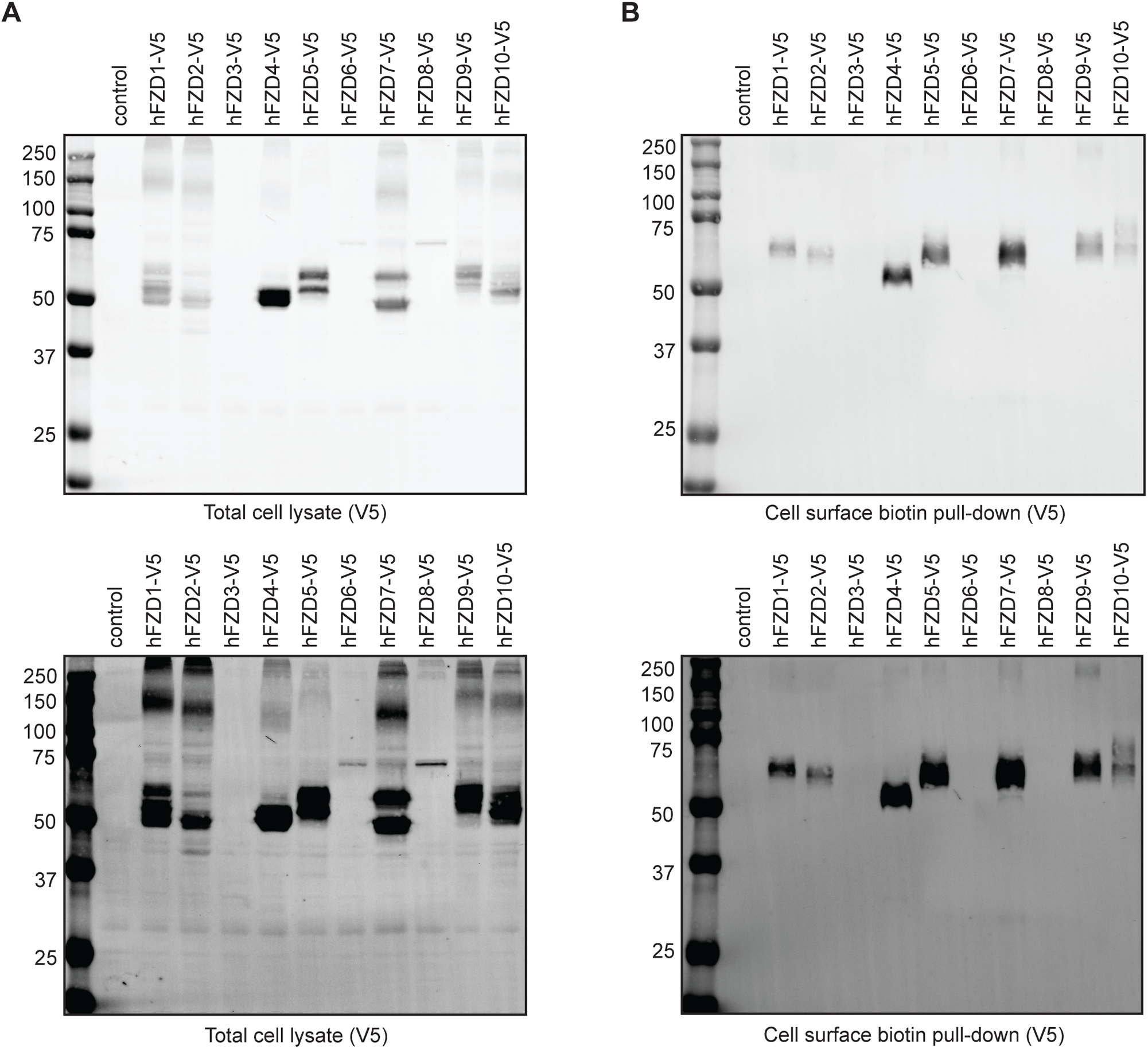
Western blot analysis of human frizzled construct distribution. **A.** Western blot analysis of V5-tagged human FZD constructs in HEK293T R/ZdKO cells in total cell lysates. **B.** Western blot analysis of the cell surface proteome of the same samples as in (**A**) after surface biotinylation and streptavidin pull-down.

**Figure S2.**
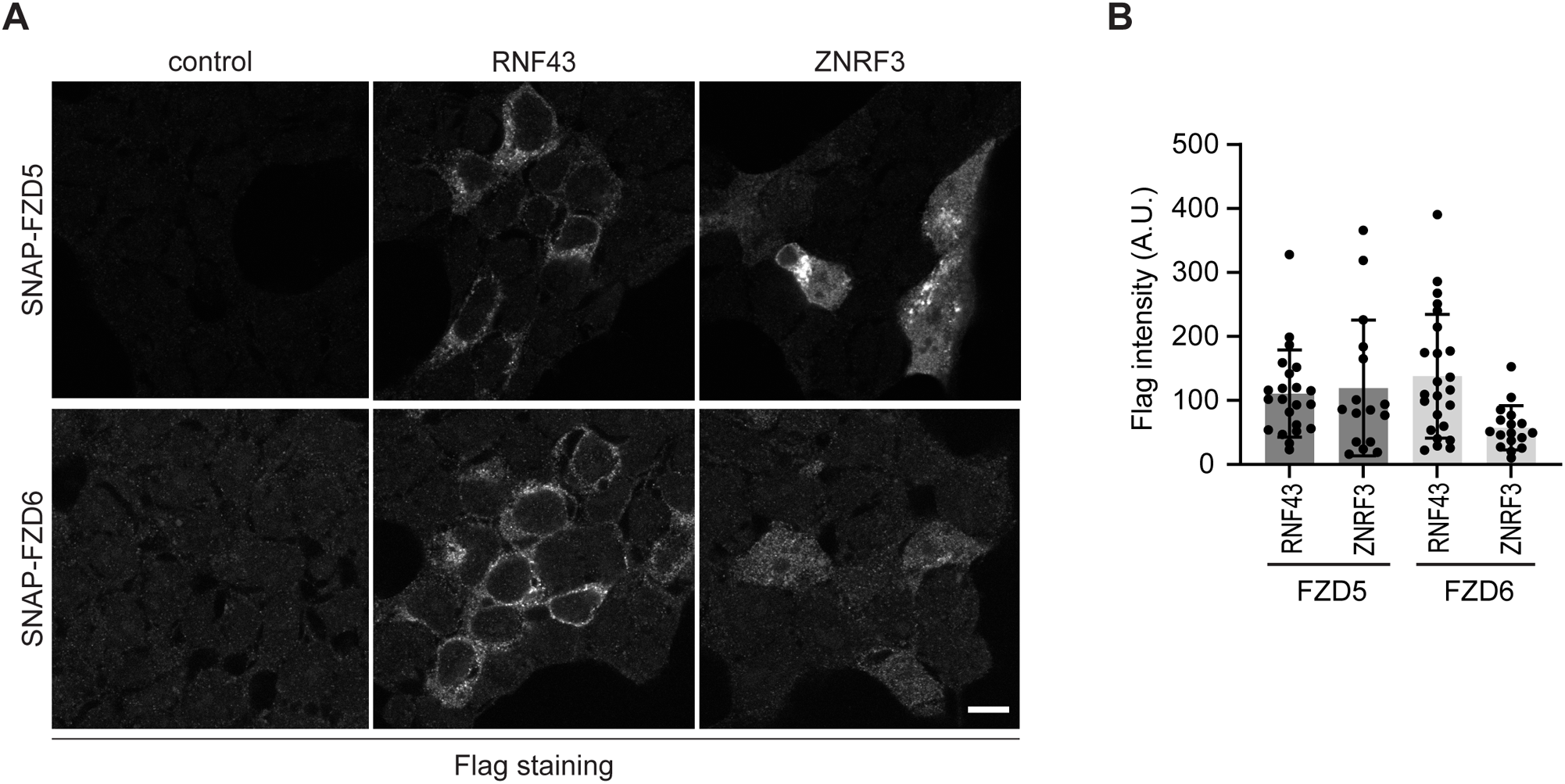
Expression levels of FLAG-tagged RNF43 and ZNRF3 in HEK293T cells. **A.** Immunofluorescence showing the subcellular localisation of Flag-tagged RNF43 and ZNRF3 of the experiment shown in figure 2D. Scale bar, 10 µm. **B**. Quantification of RNF43 and ZNRF3 expression in (A). Graph shows average Flag labelling intensities (A.U.) for the indicated conditions (n=16-24 cells).

**Figure S3.**
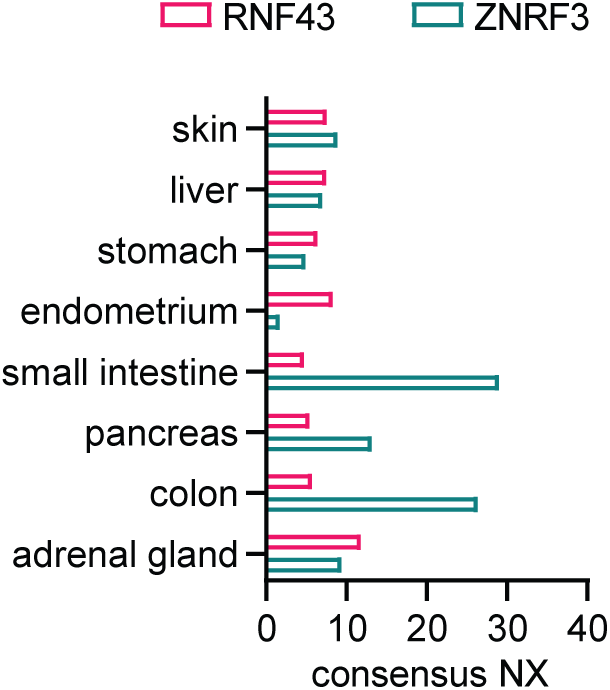
Consensus normalised expression (NX) of RNF43 and ZNRF3 among human tissues.

## Notes

### Competing Interest Statement

Madelon Maurice is cofounder and shareholder of Laigo Bio B.V. The other authors declare no competing interests.

### Summary of Updates

New experimental results added and minor textual edits.

